# ADAR-Independent A-to-I RNA Editing is Generally Adaptive for Sexual Reproduction in Fungi

**DOI:** 10.1101/059725

**Authors:** Qinhu Wang, Cong Jiang, Huiquan Liu, Jin-Rong Xu

**Affiliations:** State Key Laboratory of Crop Stress Biology for Arid Areas, College of Plant Protection, Northwest A&F University, Yangling, Shaanxi 712100, China; Department of Botany and Plant Pathology, Purdue University, West Lafayette, IN 47907, USA.

**Keywords:** RNA editing, *Fusarium graminearum*, positive selection, dN/dS, nonsynonymous

## Abstract

ADAR-mediated A-to-I RNA editing is a well-known RNA modification mechanism in metazoans that can cause nonsynonymous changes leading to amino acid substitutions. Despite a few cases that are clearly functionally important, the biological significance of most nonsynonymous editing sites in animals remains largely unknown. Recently, genome-wide A-to-I editing was found to occur mainly in the coding regions and specifically during sexual reproduction in the wheat scab fungus *Fusarium graminearum* that lacks ADAR orthologs. In this study, we found that both the frequency and editing level of nonsynonymous editing is significantly higher than those of synonymous editing, suggesting that nonsynonymous editing is generally beneficial and under positive selection in *F. graminearum*. We also showed that nonsynonymous editing favorably targets functionally more important and more conserved genes, but at less-conserved sites, indicating that the RNA editing system is adapted to fine turn protein functions by avoiding potentially deleterious editing events. Furthermore, nonsynonymous editing in *F. graminearum* was found to be under codon-specific selection and most types of codon changes tend to cause amino acid substitutions with distinct physical-chemical properties and smaller molecular weights, which likely have more profound impact on protein structures and functions. In addition, we found that the most abundant synonymous editing of leucine codons is adapted to fine turn the protein expression by increasing codon usage bias. These results clearly show that A-to-I RNA editing in fungi is generally adaptive and recoding RNA editing may play an important role in sexual development in filamentous ascomycetes.

## INTRODUCTION

RNA editing is a post-transcriptional process that alters the genome-encoded sequences in RNA transcripts by base-modifications, insertions, or deletions and may provide amino acid variations for fine-turning biological functions (Gott and Emeson 2000; Maydanovych and Beal 2006; Nishikura 2010). A-to-I RNA editing, one type of base-modification editing, is the most prevalent type of RNA editing in animals. It is mediated by adenosine deaminase acting on RNA (ADAR) enzymes, which convert adenosine (A) residue to inosine (I) via hydrolytic deamination in double-stranded RNA (dsRNA) regions of RNA molecules (Bass 2002; Nishikura 2010). Because I is interpreted as guanosine (G) by the translational machinery, A-to-I RNA editing in protein-coding regions of mRNAs may cause nonsynonymous amino acid changes (recoding). Nevertheless, editing is rarely complete and both edited and unedited versions are often expressed, which allows recoding RNA editing to create multiple protein variants from a single DNA sequence and diversify proteomes.

With the aid of high-throughput sequencing, a large number of A-to-I RNA editing sites has been identified in transcriptomes of diverse animals (Danecek et al. 2012; St Laurent et al. 2013; Chen et al. 2014; Alon et al. 2015; Picardi et al. 2015; Zhao et al. 2015). However, RNA editing in the coding regions to cause amino acid changes is generally rare in animals because the vast majority of these editing sites are located in the introns and 5’-or 3’-untranslated regions (UTRs) (Nishikura 2010; Picardi et al. 2015). In humans, over three million A-to-I editing sites have been identified but only 1,741 (approximately 0.05%) are in the coding regions (Picardi et al. 2015). In *Caenorhabditis elegans*, only 11 (0.02%) out of the 47,660 A-to-I editing sites are in the coding regions (Zhao et al. 2015). The only known exception is squid that has a total of 87,574 RNA editing sites in the coding regions, of which 57,108 (65.2%) resulting in amino acid changes (Alon et al. 2015).

Although A-to-I editing is abundant in animals, the functional significance of protein recoding RNA editing events is largely unknown. Only a small number of them have been experimentally confirmed to affect protein functions (Nishikura 2010; Pullirsch and Jantsch 2010). Amino acid changes resulting from RNA editing are known to be important for the functions of ligand-and voltage-gated ion channels and neurotransmitter receptors in invertebrates and vertebrates (Sommer et al. 1991; Seeburg 1996; Patton et al. 1997; Palladino et al. 2000). In octopus, the editing of an isoleucine (I) to valine (V) in the K+ channel has been shown to be related to temperature adaptation (Garrett and Rosenthal 2012a). However, other than these few cases that are clearly beneficial, whether most of the observed recoding RNA editing events are advantageous or not is still a point of debate or may vary among different organisms. Comparative analyses show that most recoding RNA editing sites in humans are non-adaptive and resulting from tolerable promiscuous targeting by RNA editing enzymes (Xu and Zhang 2014). However, recoding RNA editing is shown to be used for protein adaptation in invertebrates (Garrett and Rosenthal 2012b; Alon et al. 2015). Therefore, whether RNA editing is beneficial or functions as an adaptation mechanism may depend on specific organisms.

Although ADARs are unique metazoans (Jin et al. 2009; Grice and Degnan 2015) and fungi lack ADAR orthologs, recently genome-wide A-to-I RNA editing has be identified in *Fusarium graminearum* (Liu et al. 2016), a causal agent of Fusarium head blight (FHB) of wheat and barley in the US and other countries (Bai and Shaner 2004; Goswami and Kistler 2004). Among the 26,056 A-to-I editing sites that were identified specifically in sexual fruiting bodies (perithecia), 21,095 (70%) are in the coding regions, of which 78.9% resulting in amino acid changes (recoding). In total, 5,043 genes have editing sites in the coding regions. Seventy of them have premature stop codons in their ORFs that require A-to-I editing to encode full-length functional proteins, including the *PUK1* kinase gene that is important for sexual development (Wang et al. 2011 Liu et al. 2016). The editing of these premature stop codons in the ORFs is clearly essential for protein functions. However, the vast majority of editing sites in *F. graminearum* cause nonsynonymous changes and it is not clear whether they are generally advantageous or not. In this study, we analyzed the frequencies and levels of nonsynonymous editing events in *F. graminearum* and examined for their occurrences in genes of different functional importance and conserved regions of protein sequences. Results from these analyses provided unequivocal evidence that A-to-I RNA editing in fungi is generally adaptive and recoding RNA editing may play an important role in sexual development in filamentous ascomycetes.

## RESULTS

### Nonsynonymous RNA editing is generally adaptive in *F. graminearum*

Whereas synonymous editing is supposed to be neutral in general, nonsynonymous editing causing amino acids changes may be affected by selection. To determine whether the recoding RNA editing events in *F. graminearum* are adaptive, we compared the frequency of nonsynonymous editing and synonymous editing. Among the 21,095 A-to-I editing sites identified in the coding regions of *F. graminearum*, 16,985 sites are nonsynonymous and the remaining 4,110 are synonymous editing events. In the 5,043 *F. graminearum* genes with editing sites in their coding regions, a total of 1,755,425 As, if edited to G, will have nonsynonymous changes. For the rest 554,715 A sites, editing to G will be synonymous. Therefore, the actual frequency of nonsynonymous editing is 9.7 × 10^−3^ (16,985/1,755,425) and the frequency of synonymous editing is 7.4 × 10^−3^ (4,110/554,715) in *F. graminearum*. The frequency of nonsynonymous editing is over 1.3-fold higher than that of synonymous editing (*P* = 2.1 × 10^−56^, Fisher’s exacted test) (Fig. 1A), suggesting that a substantial fraction of nonsynonymous editing sites is likely under positive selection.

**Figure 1.**
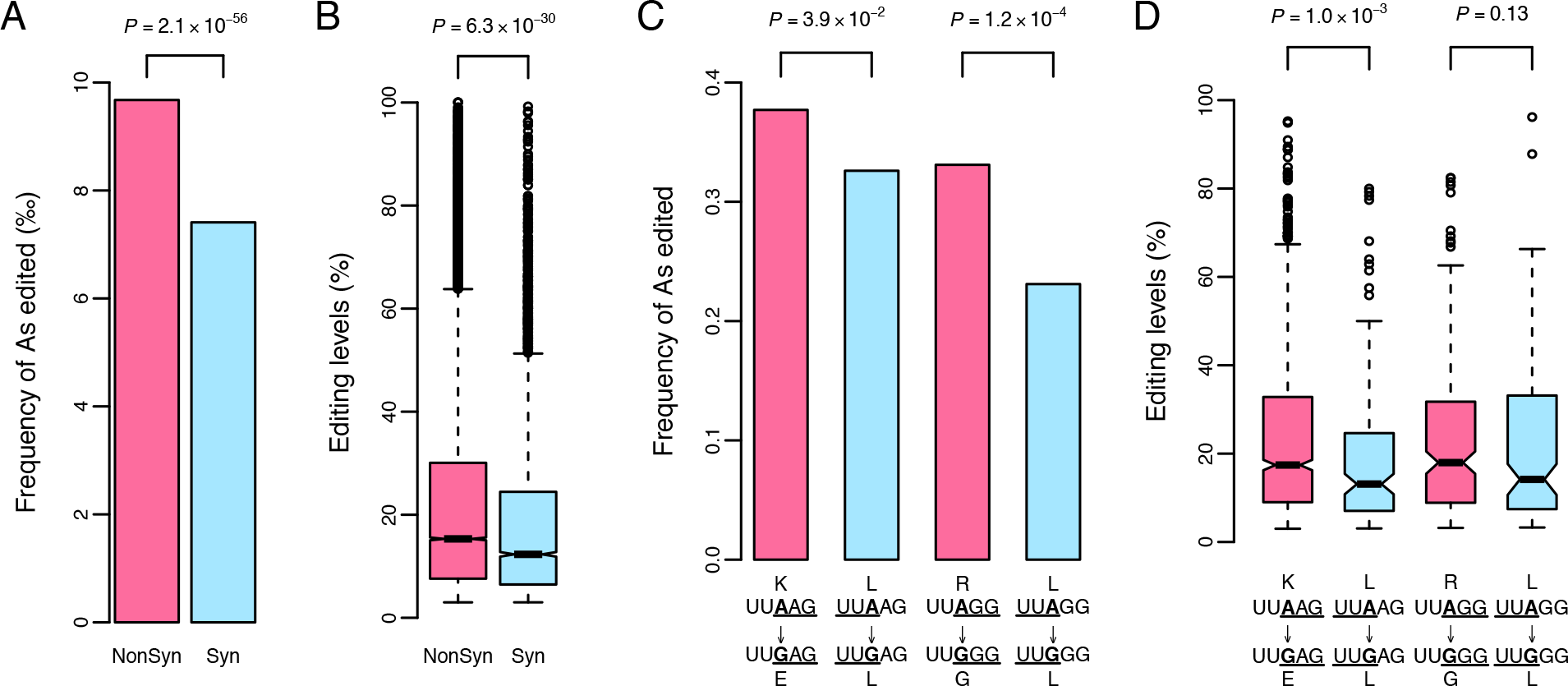
Frequencies and editing levels of nonsynonymous and synonymous A-to-I editing in *F. graminearum*. (**A**)Frequency of A sites in the coding regions of 5,043 genes underwent nonsynonymous (NonSyn) or synonymous (Syn) editing in perithecia. The *P*-value (Fishers’ exact test) labelled on the top shows that nonsynonymous editing occurs at a significantly higher frequency than synonymous editing. (**B**) Box plot of the editing levels of nonsynonymous and synonymous editing events. The *P*-value (one-tailed Wilcoxon rank sum test) labelled on the top shows that the editing level of nonsynonymous editing is significantly higher than that of synonymous editing. (**C**) Frequency of nonsynonymous and synonymous editing in the two most preferred RNA sequences UU**A**AG and UU**A**GG. The *P*-values from Fishers’ exact test are marked on the top. (**D**) Box plot of the editing levels of nonsynonymous and synonymous editing events in UU**A**AG and UU**A**GG. The *P*-values are from statistical analysis with one-tailed Wilcoxon rank sum test. In panels **C** and **D**, the upper and lower rows represent original and edited sequences, with edited As in bold and affected codons underlined.

If the observed nonsynonymous editing events are beneficial, their editing levels are expected to be higher than those of synonymous editing events because higher nonsynonymous editing levels will confer greater benefits. When editing events within the coding regions of the 5,043 genes were examined, in general, the editing levels of nonsynonymous editing sites are significantly higher than those of synonymous editing sites (Fig. 1B). Therefore, nonsynonymous editing appears to be beneficial in *F. graminearum*.

In *F. graminearum*, A-to-I editing has sequence preference at the neighboring nucleotides (Liu et al. 2016). To avoid biases due to editing preferences, we compared the frequencies of nonsynonymous and synonymous editing events with the same neighboring nucleotides in the two most favorable editing sequences UUAAG or UUAGG in the coding regions. In both sequences, editing of the middle A (bold) to G will result in synonymous or non-synonymous changes depending on the reading frames (Fig. 1C). If UUA is the leucine (L) codon, editing to UUG is synonymous. If AAG (K) or AGG (R) is the codon, editing to GAG (E) or GGG (G) will cause nonsynonymous change. For the UUAAG sequence, the K^AAG^ to R^GAG^ nonsynonymous editing has a higher editing frequency (Fig. 1C) and editing level (Fig. 1D) than the L^UUA^ to L^UUG^ synonymous editing in *F. graminearum*. For the UUAGG sequence, the frequency of nonsynonymous R^AGG^ to G^GGG^ editing is also significantly higher than that of synonymous L^UUA^ to L^UUG^ editing (Fig. 1C) although their editing levels are not statistically different (Fig. 1D). These results further indicate that nonsynonymous editing is adaptive in *F. graminearum*.

### The stop-lost editing is adaptive

Among the nonsynonymous editing sites, 323 are stop-lost editing events in which the UAG stop codon is edited to UGG tryptophan (W) codon (Liu et al. 2016). Among the synonymous editing sites, 175 are stop-retained editing events with the UAA to UGA stop codon change (Liu et al. 2016). The stop-lost editing causes the addition of an extra stretch of peptides to the C-terminal end of encoding proteins. If the observed stop-lost editing is adaptive, we expect that the frequency of UAG to UGG stop-lost editing will be higher than that of the UAA to UGA stop-retained editing. Among the 5,043 *F. graminearum* genes edited, 1,559 and 1,970 have the UAG and UAA stop codons, respectively. As predicted, the frequency of UAG being edited to UGG is 20.7% (323/1,559), which is more than two folds higher than 8.9% (175/1,970) of UAA being edited to UGA (*P* = 1.5 × 10^−23^, Fisher’s e×acted test) (Fig. 2A). Moreover, the editing levels of UAG to UGG stop-lost editing are significantly higher than those of UAA to UGA stop-retained editing (Fig. 2B), indicating that stop-lost editing is also adaptive in *F. graminearum*.

**Figure 2.**
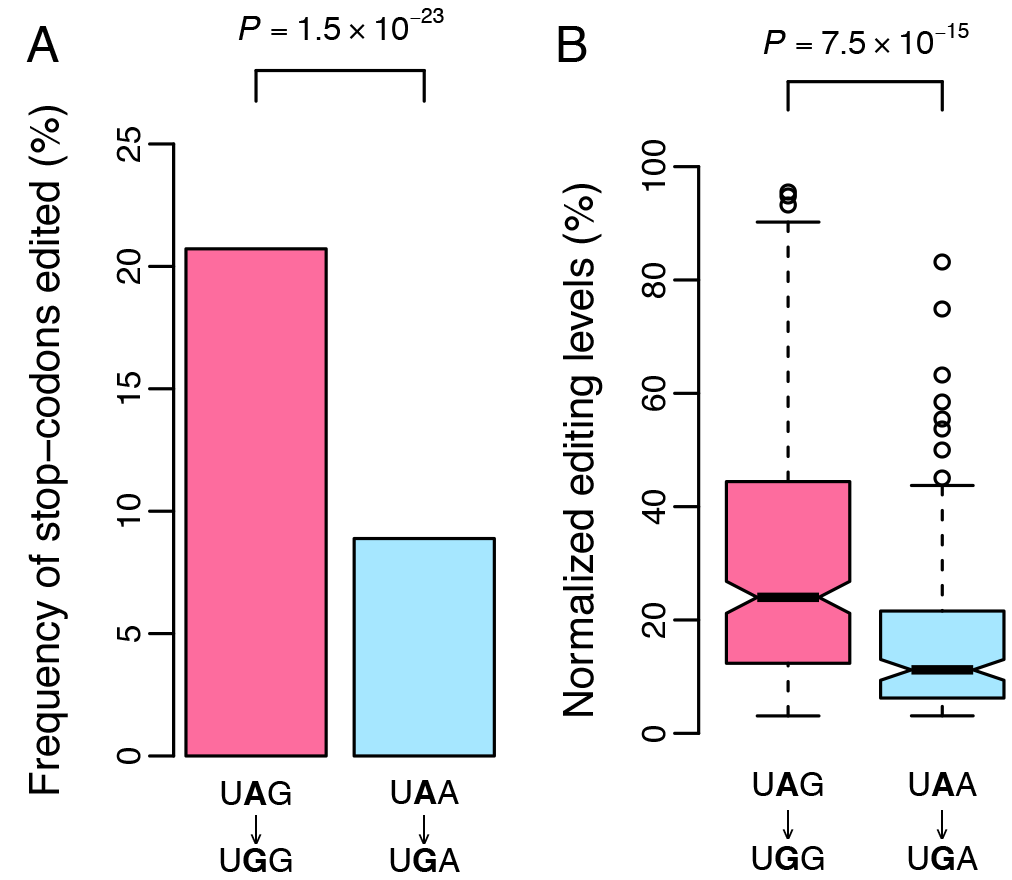
Frequencies and editing levels of UAG and UAA stop-codon editing. (**A**)The percentage of UAG and UAA stop-codons at the end of ORFs with the middle A (in bold) edited to G. Nonsynonymous editing of UAG to UGG occurs much more frequently than synonymous editing of UAA to UGA. *P*-value is from Fishers’ exact test. (**B**) Box plot of normalized editing levels of A-to-I editing in the UAG and UAA stop-codons (edited As in bold). Because the editing levels of editing events at UAG triplets is significantly higher than that of editing events at UAA triplets (Liu et al. 2016), we normalized the editing levels of UAG and UAA stop-codon editing by the median editing levels of all editing events targeted on UAG and UAA triplets, respectively. *P*-value is from statistical analysis with one-tailed Wilcoxon rank sum test.

### Nonsynonymous RNA editing is enhanced in functionally more important genes

To determine whether nonsynonymous editing is associated with functional importance of genes, we categorized the predicted *F. graminearum* genes into the essential (functionally more important) and nonessential (functionally less important) groups based on whether their orthologs are essential in the budding yeast *Saccharomyces cerevisiae* and examined for the occurrence of A-to-I editing. Among the 5,043 genes edited, 556 (11.0%) of them are in the essential group (Fig. 3A). In contrast, only 218 (2.4%) of the unedited genes belong to the essential group. Therefore, edited genes in *F. graminearum* are significantly enriched for genes that are likely functionally important (Fig. 3A).

**Figure 3.**
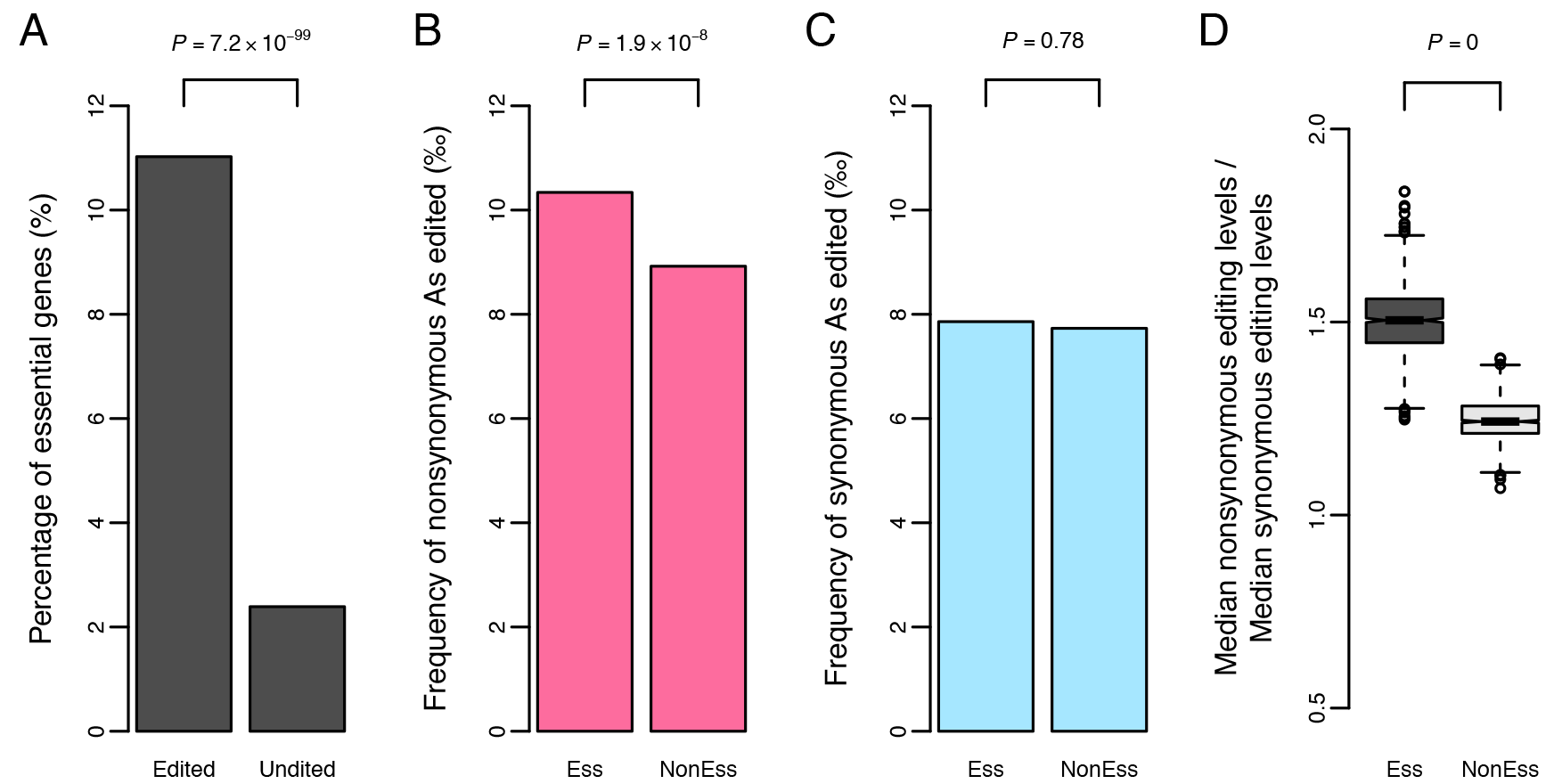
Comparative analysis of editing events in essential and nonessential genes in *F. graminearum*. (**A**)The percentage of essential (Ess) genes inferred from their yeast orthologs in edited and unedited genes in *F. graminearum*. (**B**) Frequency of nonsynonymous editing sites in essential and nonessential (NonEss) genes. (**C**) Frequency of synonymous editing sites in essential and nonessential genes. (**D**) The ratio of the median editing levels of nonsynonymous and synonymous editing sites in essential and nonessential genes. The *P*-values are from statistical analyses with Fishers’ exact test (**A-C**) and one-tailed bootstrap test with 1,000 samples (**D**).

For *F. graminearum* genes with A-to-I editing in the coding regions, the frequency of nonsynonymous editing is significantly higher in the essential gene group than in the nonessential gene group (Fig. 3B). In contrast, there is no difference in the frequency of synonymous editing between the essential and nonessential gene groups (Fig. 3C). Furthermore, the ratio between the median nonsynonymous editing level and the median synonymous editing level is significantly higher for essential genes than for nonessential genes (Fig. 3D). These results indicate that adaptive selection may make nonsynonymous RNA editing favorably targeting functionally more important genes in *F. graminearum*.

### Nonsynonymous RNA editing favorably targets genes under stronger functional constraints

To determine whether nonsynonymous editing is associated with functional constrains of genes, we identified one-to-one orthologs of genes with RNA editing events in *F. graminearum* in two closely related *Fusarium* species *F. verticillioides* and *F. solani*, and classified them into different categories based on the ratio of nonsynonymous substitution rate (dN) to synonymous substitution rate (dS) that is negatively correlated with functional constraints (Nei and Kumar 2000). Whereas the low dN/dS group contains genes with high functional constraints, genes belonging to the high dN/dS group have low functional constraints. When examined for the A-to-I editing sites, the low and high dN/dS groups have similar frequencies of synonymous editing (Fig. 4A). However, the low dN/dS group has a higher frequency of nonsynonymous editing than the high dN/dS group (Fig. 4A). These results suggest that the frequency of nonsynonymous editing increases when the dN/dS ratio decreases. In contrast, the low dN/dS group has lower editing levels than the high dN/dS group for both nonsynonymous and synonymous editing (Fig. 4B), suggesting that the editing level is restricted by the level of functional constraint.

**Figure 4.**
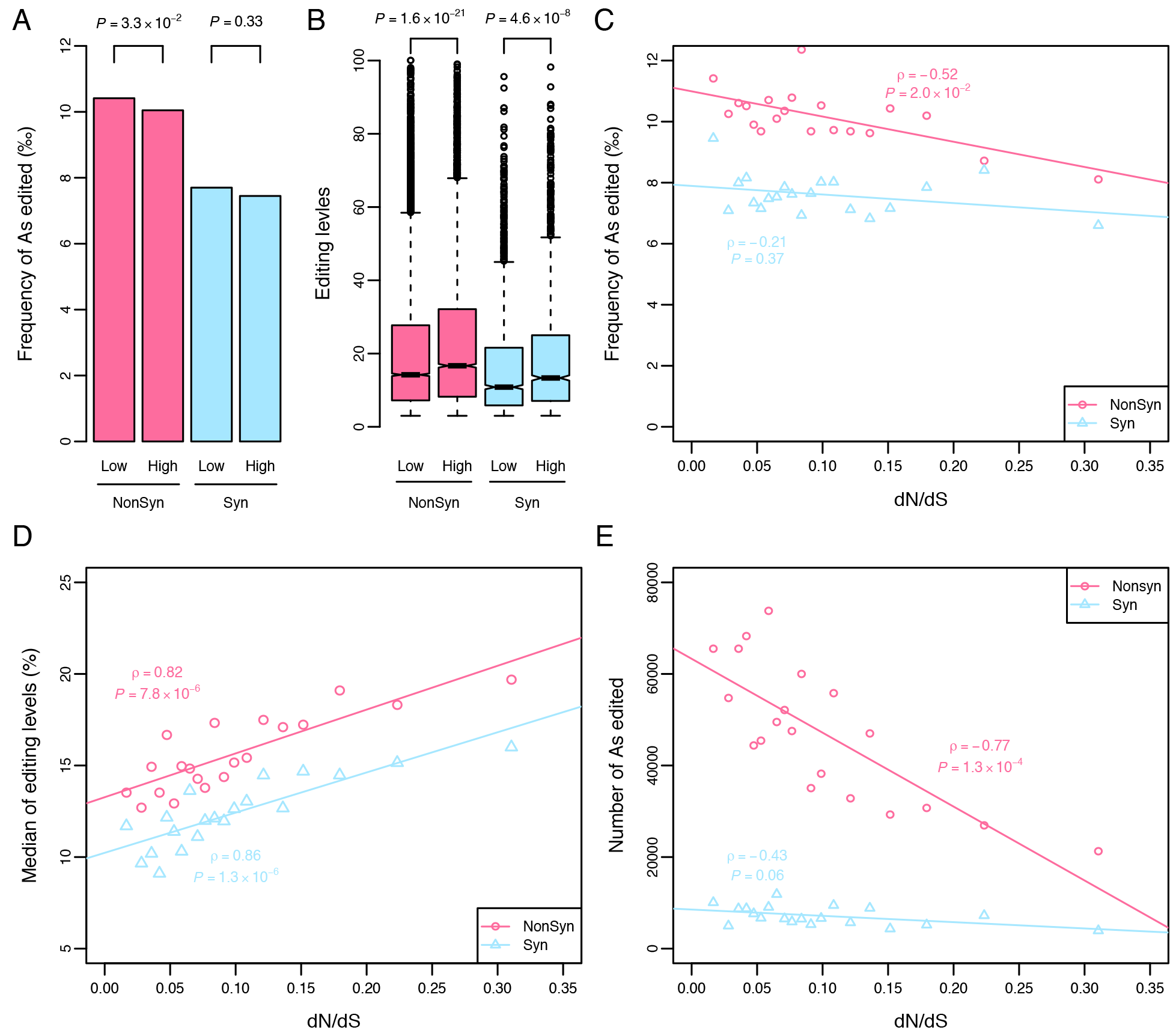
Increased A-to-I editing in *F. graminearum* genes with low ratio of nonsynonymous to synonymous rate (dN/dS) (**A**) Frequency of nonsynonymous (NonSyn) and synonymous (Syn) editing sites in the edited genes in the low or high dN/dS group. Each group represents 50% of the edited genes assigned by rank of dN/dS value calculated from *F. graminearum, F. verticillioides*, and *F. solani*. *P*-values are from Fishers’ exact tests. (**B**) Editing levels of nonsynonymous and synonymous editing sites in the edited genes in the low or high dN/dS group. *P*-value is from statistical analysis with two-tailed Wilcoxon rank sum test. (**C**) Correlation between the frequencies of nonsynonymous or synonymous editing sites and the medians of dN/dS ratios for each group. (**D**) Correlation between the median editing levels of nonsynonymous or synonymous editing sites and the medians of dN/dS ratios for each group. (**E**) Correlation between the editing intensity (the total number of As edited) and the medians of dN/dS ratios for each group. In **C-E**, each point represents a group with 5% of edited genes. Rho and *P*-value are from statistical analyses with two-tailed Spearman’s rank correlation test.

We then divided the edited genes into 20 groups with equal number of genes in each group based on the rank of dN/dS values and examined for the occurrence of RNA editing. A significant negative correlation between the frequency of nonsynonymous editing and dN/dS values was identified (Fig. 4C). In contrast, there is no significant correlation between the frequency of synonymous editing and dN/dS values (Fig. 4C). As a control, we generated the same number of editable A sites by randomly selecting the A sites with U at the −1 position from the coding regions of the 5,043 edited genes, which accounts for the observed strong sequence preference of U at the −1 position of edited sites in *F. graminearum* (Liu et al. 2016). The frequency of random nonsynonymous or synonymous editable sites had no significant correlation with the dN/dS values (Fig. S1). Furthermore, although the medians of editing levels of both nonsynonymous and synonymous editing are positively correlated with dN/dS values (Fig. 4D), the intensity of A-to-I editing measured by the number of total edited As also has a significant negative correlation with dN/dS values for nonsynonymous editing but not for synonymous editing (Fig. 4E). These results further indicate that adaptive selection may affect nonsynonymous RNA editing to target genes under stronger functional constraints.

### Nonsynonymous RNA editing favors less-conserved positions

To evaluate the evolutionary features of edited sites, we generated codon alignments for each ortholog family among 14 Sordariomycete fungi (Supplementary Fig. S2) and compared the conservation of the edited A sites and affected amino acid residues in *F. graminearum* as measured by Shannon entropy (Shannon 1948). The lower the Shannon entropy, the higher the evolutionary conservation. Compared to the random editable sites mentioned above, the Shannon entropy is significant higher at the nucleotide (Fig. 5A) or amino acid (Fig. 5B) level for nonsynonymous editing sites but not for synonymous sites. The elevated Shannon entropy for synonymous editing sites (Fig. 5A) is consistent with the fact that nucleotide substitutions at the third position of a codon are generally synonymous and tend to occur more frequently. Therefore, nonsynonymous edited A sites in *F. graminearum* tend to recode the less-conserved nucleoside As at less conserved amino acid residues.

We then examined for the association of editing levels with the conservation of edited A sites. The editing sites with higher editing levels have higher Shannon entropy (lower conservation) at the nucleotide (Fig. 5C) and amino acid (Fig. 5D) levels for nonsynonymous editing sites but not for synonymous sites. These results indicate that nonsynonymous RNA editing in fungi favors less-conserved coding sites to avoid potentially detrimental effects on protein functions.

**Figure 5.**
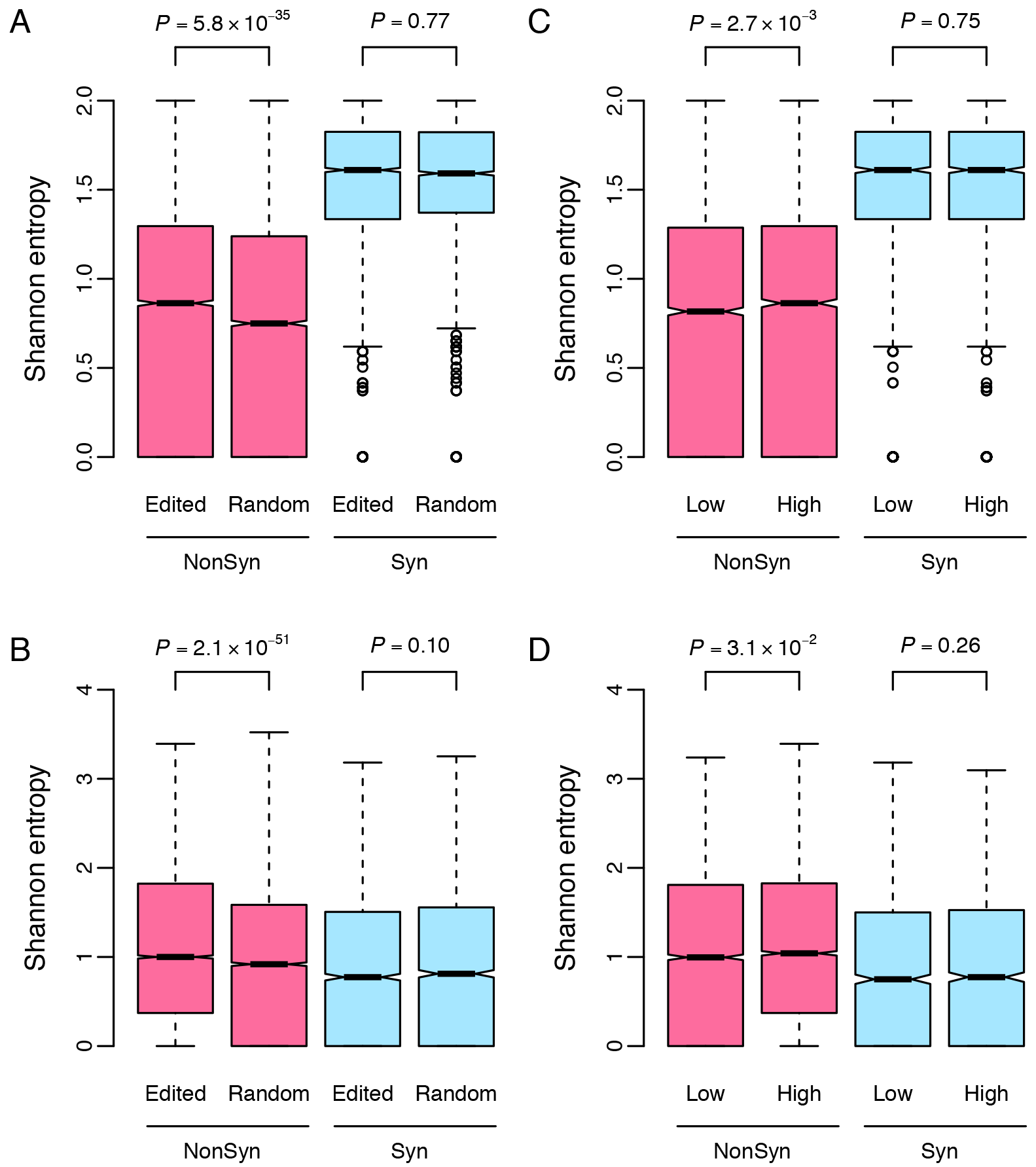
Conservation of edited nucleotide and amino acid sites in *F. graminearum*. Box plot comparison of the Shannon entropy of nonsynonymous (NonSyn) and synonymous (Syn) edited sites with those of random editable sites in the alignments of one-to-one orthologs from 14 Sordariomycetes at nucleotide level (**A**) and amino acid level (**B**). The random editable sites are generated by randomly selecting 21,095 editable A sites with U at the −1 position from the coding regions of the 5,043 edited genes. Box plot comparison of the Shannon entropy of editing positions grouped by low or high editing level for nonsynonymous and synonymous editing at nucleotide level (**C**) and amino acid level (**D**). Each group represents 50% of the edited genes assigned by rank of editing levels. Because Shannon entropy measures the uncertainty of the editing sites, lower value indicates higher conservation, and vice versa. *P*-values in **A-D** are from statistical analyses with two-tailed Wilcoxon rank sum test.

### Nonsynonymous RNA editing is under codon-specific selection

To determine whether RNA editing is under codon-or residue-specific selection, we compared the frequency of observed codon or residue changes to what is expected by chance, accounting for the observed strong sequence preference for U at the −1 position of edited sites and number of editing events. For most types of amino acid substitutions that caused by nonsynonymous A-to-I editing, the observed editing frequencies are significantly different from what is expected for a random selection (Fig. 6A). Residue changes rendered by RNA editing with more significantly higher frequency than expected tend to cause more drastic amino acid changes. For example, the top four most frequent types of residue changes, lysine (K) to glutamic acid (E), asparagine (N) to aspartic acid (D), serine (S) to glycine (G), and arginine (R) to glycine (G) are at least two-times more frequent than expected, and result in a drastic difference in the physical-chemical properties of amino acid residues (Fig. 6A). It is likely that most of the nonsynonymous editing events causing these types of residue changes are beneficial and fixed by positive selection. Nevertheless, the tyrosine (Y) to cysteine (C), threonine (T) to alanine (A), and isoleucine (I) to valine (V) changes are far less frequent than expected (Fig. 6A). Editing events resulting in these amino acid changes may be detrimental to the protein functions of edited genes and are eliminated by purifying selection. Therefore, it is likely that nonsynonymous RNA editing is under codon-or residue-specific selection in *F. graminearum*.

**Figure 6.**
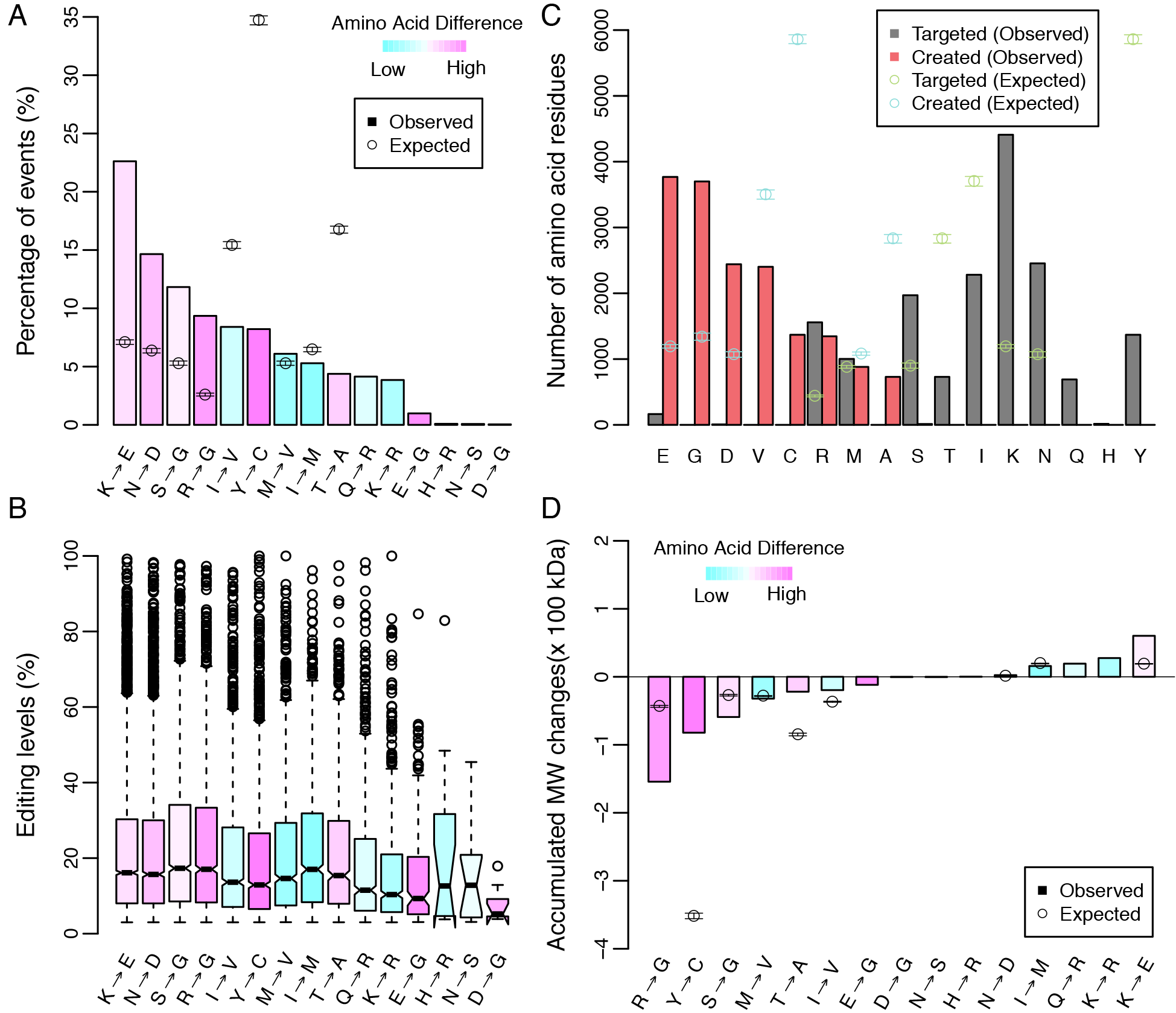
Amino acid or codon changes caused by A-to-I RNA editing in *F. graminearum*. (**A**) Distribution of amino acid or codon changes caused by A-to-I editing. (**B**) Editing levels for different types of amino acid or codon changes caused by A-to-I editing. (**C**) The number of amino acids targeted and created by A-to-I editing. (**D**) Effects of A-to-I editing on the molecular weight (MW) of proteome. The accumulated molecular weight changes represent the sum of all molecular weight changes caused by editing events for each type of amino acid substitution. In **A**, **B**, and **D**, Grantham scores (Grantham 1974) of nonsynonymous amino acid changes were mapped from cyan to magenta to represent the amino acid difference between the residue targeted and created, from low to high. The higher Grantham score, the greater difference in physical-chemical properties between two amino acids. In **A**, **C**, and **D**, the expected values (circles with error bars) were calculated from the 21,095 random editable A sites with U at the −1 position randomly selected from the coding regions of the 5,043 edited genes. The expected value of 0 is not plotted. Error bars represent the standard deviations from 1,000 times sampling. For each type of amino acid substitution with the expected value of non-zero, the observation is significantly different from what is expected by two-tailed one sample *t*-test.

When the editing levels of codon change editing sites were compared (Fig. 6B), we found that the favored codon changes may be under positive selection (such as S to G and R to G editing) tend to have higher editing levels than the disfavored codon changes (such as Y to C and I to V editing) (Fig. 6B). These results further indicate that selection on nonsynonymous editing is codon or residue-specific in filamentous fungi.

### Nonsynonymous RNA editing shifts the composition and molecular weight of proteome

By comparing the predicted and observed numbers of codons targeted by nonsynonymous editing, we found that the K, N, S, and R residues were targeted for RNA editing over two times more frequently than expected. In contrast, RNA editing occurred at the Y, T, and I residues is far less frequently than expected (Fig. 6C). On the other hand, among the amino acid residues created by A-to-I editing, the creation of the C, A, and V residues occurs less than expected. However, the E, G, and D residues are created by RNA editing at a level significantly higher than the expected frequency (Fig. 6C), indicating that RNA editing favorably creates these residues. Among all the 20 amino acid residues, only the R and M residues have similar numbers of being targeted and created by RNA editing (Fig. 6C). The difference between amino acid residues targeted for editing and residues created by RNA editing indicate that nonsynonymous editing tends to change the overall amino acid composition of proteins in *F. graminearum*.

Furthermore, we examine the effect of RNA editing on the molecular weight of proteins by calculating the accumulated molecular weight changes of the amino acid residues created relative to targeted. Overall, most of the residue changes caused by A-to-I editing result in a reduction in the molecular weights of encoding proteins (Fig. 6D). The accumulative effects of codon change editing events reduce the molecular weights of proteins encoded by edited genes. These observations indicate that nonsynonymous editing may affect the amino acid compositions and molecular weights of the proteome in *F. graminearum*.

### Synonymous editing increases codon bias

Although synonymous editing is thought to be neutral, we observed that the editing events resulting in L to L synonymous changes occur more frequently than expected, whereas the other types of synonymous changes are not (Fig. 7A). Their editing levels are also relatively higher than those of the other types of synonymous changes (Fig. 7B). Interestingly, both CU**A** (L) to CU**G** (L) and UU**A** (L) to UU**G** (L) changes convert a less frequently used leucine codon to a more frequently used one (Fig. 7C). We therefore defined codon bias change score (see Methods) to measure the total contribution of each type of synonymous changes to the codon bias. The CU**A** (L) to CU**G** (L) and UU**A** (L) to UU**G** (L) codon changes are not only the most frequent synonymous editing events but also have important roles in increasing codon bias in the *F. graminearum* proteome (Fig. 7C). Therefore, it is likely that the L to L synonymous editing also are under positive selection to fine turn the expression of proteins by increasing the codon usage bias.

**Figure 7.**
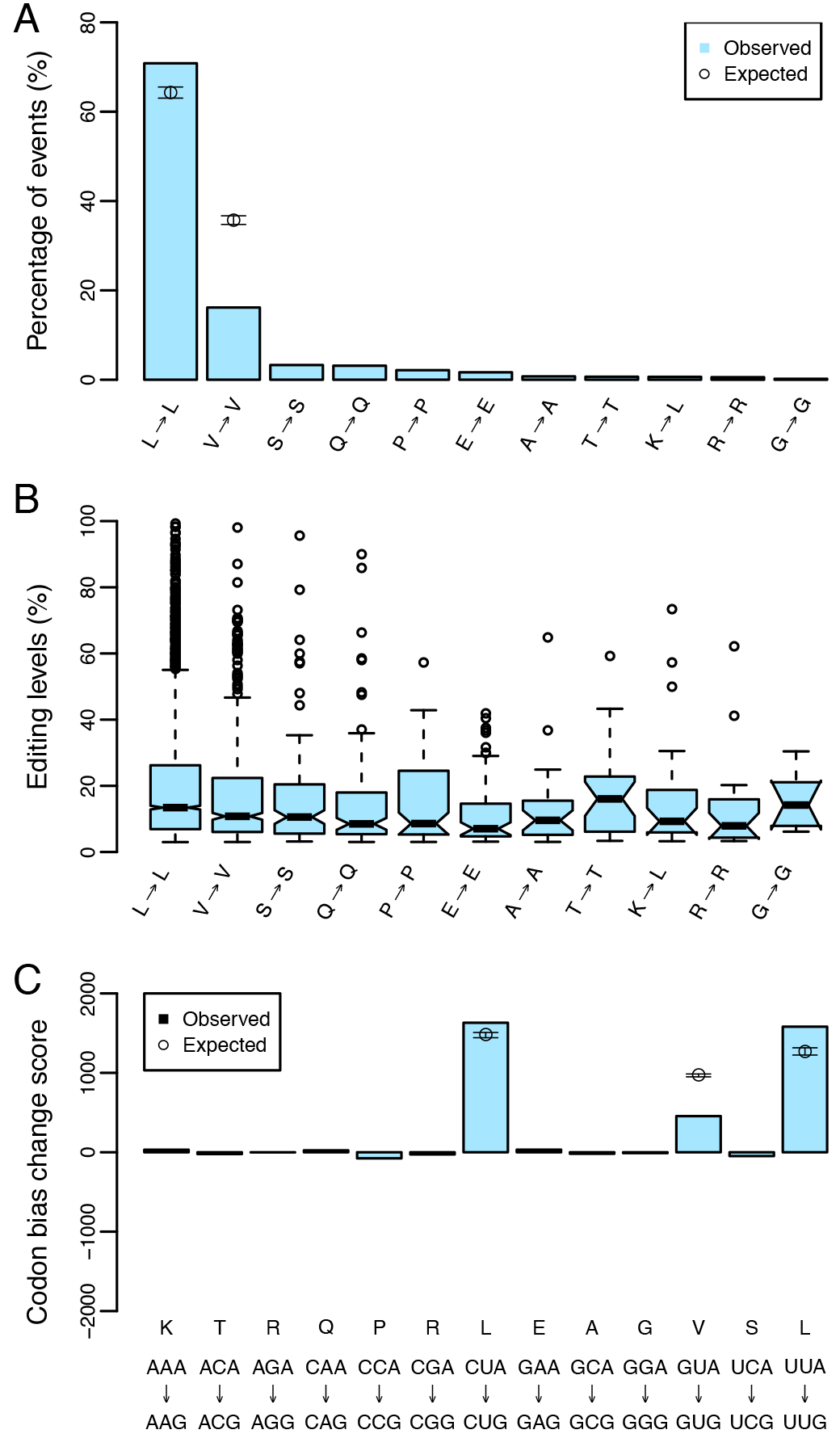
Changes of codon and codon bias in synonymous editing. (**A**) Distribution of synonymous codon changes caused by A-to-I editing. (**B**) Editing levels for different types of synonymous codon changes caused by A-to-I editing. (**C**) Codon bias change caused by synonymous editing. For each synonymous codon substitution, the codon bias change score measuring the total changes of codon priority caused by synonymous editing. The expected values (circles with error bars) were calculated from the 21,095 random editable A sites with U at the −1 position randomly selected from the coding regions of the 5,043 edited genes. The expected value of 0 is not plotted. Error bars represent the standard deviations from 1,000 times sampling. For L to L and V to V amino acid substitutions, the observation is significantly different from what is expected by two-tailed one sample *t*-test.

## DISCUSSION

Despite a few cases of recoding RNA editing are known to be functionally important (Nishikura 2010; Pullirsch and Jantsch 2010), the biological significance of nonsynonymous editing events in animals remains an open question. In humans, A-to-I editing sites identified in the coding regions are largely deleterious rather than beneficial, as both the frequency and level of nonsynonymous editing events are significantly lower than those of synonymous editing events (xu and Zhang 2014). Furthermore, nonsynonymous RNA editing is rarer in essential genes and genes under stronger functional constraints (xu and Zhang 2014). In *F. graminearum*, however, the frequency and level of nonsynonymous editing events are all significantly higher than those of synonymous editing events. Furthermore, nonsynonymous RNA editing in *F. graminearum* favorably targets genes with more important functions and genes under stronger functional constraints. These results indicate that recoding RNA editing in *F. graminearum*, unlike those in humans and other animals, is adaptive and selection makes it fine turning the function of more important and more conserved genes.

Nonsynonymous editing events in both human and squid are more frequently observed in less conserved positions of protein coding genes (Alon et al. 2015; Grassi et al. 2015). However, analysis of 16 A-to-I RNA editing sites in the Drosophila nervous system suggested that many of them alter highly conserved and functionally important positions in proteins (Hoopengardner et al. 2003). Nevertheless, a comparative study of Kv2 K^+^ channels from insects indicated that A-to-I RNA editing usually occurs at less-conserved positions in the highly conserved coding regions (Yang et al. 2008). In this study, we found that nonsynonymous editing in *F. graminearum* favorably alters less-conserved amino acid residues in highly conserved genes. It is likely that RNA editing is adapted to avoid the A sites that nonsynonymous editing may have deleterious effects or be detrimental to protein functions in fungi.

In *F. graminearum*, our analysis showed that nonsynonymous RNA editing is likely under codon-or residue-specific selection. Whereas the K, N, and S codons have the highest frequency of recoding editing events, the E, D, and G residues have the highest frequency of being created by nonsynonymous editing. In contrast, the Y, T, and I residues are least favorable targets for recoding editing and creation of C, A, and V residues are far less frequently than expected. In *Drosophila*, R residues are edited far less frequently than expected (Garrett and Rosenthal 2012b). In *F. graminearum*, recoding editing events at the R residues occurs at a higher frequency than expected. However, the overall number of R residues edited is similar to the number of R residues created by A-to-I editing in *F. graminearum*. In cephalopods, I residues are targeted for editing approximately three-fold more frequently than expected (Garrett and Rosenthal 2012b), which is also different from I being one of the least favorable targets for recoding editing in *F. graminearum*. In both human and fly, nonsynonymous RNA editing tends to avoid drastic amino acid changes. Nonsynonymous events leading to similar amino acid changes are more frequent than those causing drastic changes in the physical-chemical properties of amino acids (Grassi et al. 2015). In *F. graminearum*, however, nonsynonymous events resulting in changes to amino acid residues of different physicochemical properties are more frequent than expected. Therefore, unlike in metazoans, A-to-I RNA editing in filamentous fungi may play an important role in the diversification of protein functions.

Although it has tissue or developmental stage preference in animals, A-to-I RNA editing has been identified in virtually all the tissues examined (Picardi et al. 2015). In contrast, A-to-I editing is stage-specific and occurs only during sexual reproduction in *F. graminearum* (Liu et al. 2016). Intriguingly, fungal nonsynonymous RNA editing favors genes under stronger functional constraints (lower dN/dS value). The greater the functional constraint, the higher frequency of being edited. Sexual reproduction plays a critical role in the Fusarium head blight (FHB) disease cycle because forcibly discharged ascospores (sexual spores) from overwintering perithecia serve as the primary inoculum (Schmale III et al. 2005; Trail 2009). Any molecular mechanisms that increase genetic variability during sexual reproduction may drive the adaptation of *F. graminearum* for acclimatization during sexual development and subsequent infection. However, genetic recombination (Burt 2000) and spontaneous mutation during meiosis (Koltin et al. 1975; Rattray et al. 2015) are permanent and hardwired, which restricts the extent of genetic variations at functionally critical amino acid residues that may be under purifying selection. Thus, RNA editing during sexual reproduction may serve as one strong driving force for adaptive evolution as proposed (Gommans et al. 2009). Complementary to genomic mutations, RNA editing may provide sequence variations in highly conserved genes that are not accessible for genetic mutation in the genomic sequence because of their functional constraint.

RNA editing is seldom complete with editing levels at a specific site ranging from a few to almost 100%. Some editing events, such as the editing events in *PUK1* and other pseudogenes (Liu et al. 2016), have relative high editing levels. These editing events may cause adaptively important protein variants and thereby are under directional selection to increase their editing levels. On the other hand, many editing events have relative low editing levels and may lack obviously beneficial effects under normal conditions. The new variants created by these RNA editing events could be maintained at a low fraction by balancing selection to potentially increase the genetic variability of organisms. For example, incomplete recoding events at three different sites in a gene will theoretically generate 2^3^ = 8 different protein variants. The remarkable variations could allow for fast acclimation after environmental change and facilitate adaptive evolution. Moreover, besides increasing variations, nonsynonymous events more frequently lead to more drastic changes in the physical-chemical properties and molecular weights of amino acids in *F. graminearum*. Recent studies have showed that RNA editing can respond to acute temperature changes and be used for temperature adaptation by increasing the flexibility of protein (Garrett and Rosenthal 2012a; Garrett and Rosenthal 2012b; Savva et al. 2012). The large number of recoding events during sexual reproduction may provide greater flexibility of proteins for cold adaptation and responding to other environmental variables in fungal pathogens.

## METHODS

### Classification of the essential and nonessential genes in *F. graminearum*

The orthologous genes between *S. cerevisiae* and *F. graminearum* were identified by using OrthoMCL (Li et al. 2003) with default parameters. The essential and nonessential yeast genes were obtained from the *Saccharomyces* genome database (http://www.yeastgenome.org). A total of 3,419 *F. graminearum* genes have orthologs in yeast. Of these, 774 genes are orthologous to the genes essential for viability in yeast, which are considered to be functional more important genes (essential genes) in this study. There are 2,636 genes orthologous to the genes nonessential for viability in yeast, which are considered to be less important (nonessential genes) in *F. graminearum*. In the 5,043 genes with editing events in their coding regions, 556 and 1,488 genes were classified to be essential and nonessential genes, respectively.

### Calculation of the dN/dS

To calculate the *dN* (nonsynonymous substitution rate) / *dS* (synonymous substitution rate) ratio, orthologs of the 5,043 edited *F. graminearum* genes in *F. verticillioides* and *F. solani* were identified by OrthoMCL (Li et al. 2003) with default parameters. Of these, 4,101 genes have one-to-one orthologs in all three fungi. For the orthologous genes, their protein sequences were aligned with MUSCLE (Edgar 2004) and codon alignments were generated with PAL2NAL (Suyama et al. 2006). The *dN/dS* ratios were calculated by yn00 implemented in the PAML package (Yang 2007).

### Analysis of the conservation of editing sites

For each edited gene in *F. graminearum*, codon alignments of orthologous genes from 14 Sordariomycetes (Supplementary Fig. S2) were generated as above description. Shannon entropy (Shannon 1948) was used to measure the conservation of the corresponding nucleotide and amino acid sites that are targeted by A-to-I editing in the alignment. The definition of Shannon entropy for such a targeted site is:***H(X)*** = − Σ *x_i_* log_2_ *x_i_* where *X* is a series of frequencies of nucleotides or amino acids at the editing sites with possible value {*x*_1_,…,*x_n_*], for nucleotide sequence, *n* = 4, and for amino acid sequence, *n* = 20. Lower Shannon entropy indicates a higher degree of conservation.

### Analysis of codon substitutions

The observed codon substitutions were counted with the A-to-I RNA editing data of *F. graminearum* (Liu et al. 2016). The expected codon substitutions were estimated with 1,000 random editable datasets, each of which was generated by randomly selecting 21,095 editable A sites with U at the −1 position from the coding regions of the 5,043 *F. graminearum* edited genes. Changes in molecular weight for each type of amino acid substitution were calculated by the following formula: *ΔM* = Σ(*M_c_* − *M_t_*), where *M_c_* and *M_t_* represent the molecular weight of amino acid residues created and targeted, respectively. The codon bias change score for each type of synonymous codon substitution is defined as: *S* = Δlog *f_c_/f_t_*, where *f_c_* and *f_t_* represent the frequencies of codon usage for the codons created and target by synonymous editing, respectively. For each nonsynonymous editing, the difference between amino acid residues is measured and illustrated according to the ranks of Grantham scores (Grantham 1974). The Grantham score predicts the difference in the physical-chemical properties of the amino acid substitutions. The higher Grantham score, the greater difference in physical-chemical properties between two amino acids.

### Statistical analysis

All statistical analyses were performed using *R* (https://www.r-project.org). Five statistical methods, *t*-test, Fisher’s exacted test, Wilcoxon rank sum test, Spearman’s rank correlation test, and Bootstrap were used.

## ACKNOWLEDGEMENTS

We thank Dr. Chenfang Wang for fruitful discussions. This work was supported by grants from the Northwest A & F University Young Talent program (for Huiquan Liu), the National Basic Research Program of China (2012CB114002 and 2013CB127703), and US Wheat and Barley Scab Initiative.

